# Vimentin intermediate filaments mediate cell shape on visco-elastic substrates

**DOI:** 10.1101/2020.09.07.286237

**Authors:** Maxx Swoger, Sarthak Gupta, Elisabeth E. Charrier, Michael Bates, Heidi Hehnly, Alison E. Patteson

## Abstract

The ability of cells to take and change shape is a fundamental feature underlying development, wound repair, and tissue maintenance. Central to this process is physical and signaling interactions between the three cytoskeletal polymeric networks: F-actin, microtubules, and intermediate filaments (IFs). Vimentin is an IF protein that is essential to the mechanical resilience of cells and regulates cross-talk amongst the cytoskeleton, but its role in how cells sense and respond to the surrounding extracellular matrix is largely unclear. To investigate vimentin’s role in substrate sensing, we designed polyacrylamide hydrogels that mimic the elastic and viscoelastic nature of in vivo tissues. Using wild-type and vimentin-null mouse embryonic fibroblasts, we show that vimentin enhances cell spreading on viscoelastic substrates, even though it has little effect in the limit of purely elastic substrates. Our results provide compelling evidence that the vimentin cytoskeletal network is a physical modulator of how cells sense and respond to mechanical properties of their extracellular environment.

## I. INTRODUCTION

The physical properties of the extracellular environment impact many cellular and tissue-level functions [1–3]. Cells sense and respond to physical forces by converting them into cellular signals through a process known as mechano-sensing [4]. Recent studies indicate that the cell cytoskeleton plays a key role in transmitting forces from cell-matrix adhesions to the nuclear surface, which can alter nuclear shape and lead to changes in gene expression [5–7]. The cell cytoskeleton is comprised of three polymeric networks: F-actin, microtubules, and intermediate filaments (IF). Vimentin is an IF protein that is essential to the structural integrity of the cell [8, 9] and maintaining nuclear shape [10], but its role in in cellular mechano-sensing is unclear.

A key mechano-sensing signature in fibroblasts is increasing cell spread area on elastic substrates of increasing stiffness [11, 12]. A recent study has shown that loss of vimentin decreases cell spreading on soft elastic substrates but increases cell spreading on stiff substrates [13]. In the range of physiological-relevant stiffnesses, on the order of 1-10 kPa in shear moduli, the spread areas of wild-type and vimentin-null mEF, however, are nearly indistinguishable [13]. On elastic substrates, the ability of cells to generate traction stresses and change shape depends on the shear modulus of the cytoskeleton, which is dominated by F-actin and microtubules [6]. Yet, most biological tissues are not only elastic but viscoelastic, capable of dissipating applied stresses on timescales relevant to cellular mechanical sensing [14–16]. Substrate viscoelasticity induces changes in cell morphology and cytoskeletal structure, reducing stress fiber formation and suppressing cellular adhesions [16]. On viscoelastic substrates, stresses imposed by a spreading cell dissipate and cell spreading is set by a balance between the stress relaxation time scales of the viscoelastic substrate and the cell focal adhesion turnover rate [17]. The role of vimentin in the mechanical resilience of cells [13, 18, 19] and its role in focal adhesion assembly [20, 21] suggest that despite the modest effect of vimentin on elastic surfaces, their effects in more physiologically-relevant viscoelastic settings might be more evident.

To assess the effect of vimentin on cell surface sensing, here we use polyacrylamide hydrogels that model the elastic and viscoelastic properties of real tissues. Using wild-type and vimentin-null mouse embryonic fibroblasts (mEF), example immunofluorescence images are show in Fig. 2, we find that unlike substrate stiffness, small changes in substrate viscoelasticity has profound effects on how vimentin impacts cell spreading. Unlike elastic substrates, loss of vimentin significantly reduces cell spread area on substrates with viscous dissipation. Our results suggest vimentin intermediate filaments are a significant contributor to cellular mechano-sensing and could drive differences in cells spreading and motility in tissue.

## II. MATERIALS AND METHODS

### A. Cell Culture

Wild-type mEFs and vimentin-null mEFs were kindly provided by J. Ericsson (Abo Akademi University, Turku, Finland) and maintained in Dulbecco’s Modification of Eagle’s Medium (DMEM) + 4.5 g/L glucose, L-glutamine, and sodium pyruvate. Culture media is supplemented with 10% fetal bovine serum, 1% penicillin streptomycin with 25 mM HEPES and 1% non-essential amino acids. Cell cultures were maintained at 37° C with 5% CO_2_. Cultures were passaged when they reached 70% confluence.

### B. Immunofluorescence

Cells were fixed for immunofluorescence using 4% paraformaldehyde (Fisher Scientific, Hampton, New Hampshire). Cells membranes are permeabilized with 0.05% Triton-X (Fisher BioReagents, Hampton, New Hampshire) in PBS for 15 minutes at room temperature and blocked with 1% bovine serum albumin (BSA) (Fisher BioReagents, Hampton, New Hampshire) for 30 minutes at room temperature.

For vimentin visualization, cells were incubated with primary anti-vimentin polyclonal chicken antibody (Novus Biologicals, Centennial, Colorado) diluted 1:200 in 1% BSA in PBS for 2 hours at room temperature; the secondary antibody anti-chicken Alexa Fluor 488 (Invitrogen Carlsbad, California) was used at a dilution of 1:1000 in 1% BSA in PBS incubated covered from light for 1 hour at room temperature. For visualizing paxillin we use primary anti-paxillin mouse antibody (BD Biosciences, San Jose, California) diluted 1:400 in 1% BSA in PBS and incubated for 2 hours at room temperature; secondary antibodies were anti-mouse Alexa Fluor 633 (Invitrogen, Carlsbad, California) at a dilution of 1:1000 in 1% BSA in PBS incubated covered from light for 1 hour at room temperature. Cells were stained for DNA using Hoechst 33342 (Molecular Probes, Eugene, Oregon) at a concentration of 1:1000 in 1% BSA in PBS; cells were stained for actin using rhodamine phalloidin 565 (Invitrogen, Carlsbad, CA) at a dilution of 1:200 in 1% BSA in PBS. Cells were incubated for 1 hour covered from light at room temperature while staining for both DNA and actin.

Cells were imaged using a Leica DMi8 (Leica, Bannockburn, Illinois) equipped with a spinning disk X-light V2 Confocal Unit using a HC PL APO 40x/1.19 W CORR CS2 water immersion objective. Images were acquired using the VisiView software (Visitron Systems, Puchheim, Germany) using a 10.0 micron z-step series with 0.20 micron steps and analyzed with ImageJ (NIH, Bethesda, Maryland).

### C. Bright field imaging and cell area measurement

Bright field imaging was performed using a Nikon Eclipse Ti (Nikon Instruments, Melville, NY) inverted microscope equipped with an Andor Technologies iXon em+ EMCCD camera (Andor Technologies, South Windsor, CT). Cells were maintained at 37° C and 5% CO_2_ using a Tokai Hit (Tokai-Hit, Shizuoka-ken, Japan) stage top incubator and imaged used a 10x objective. Cell areas were traced manually using ImageJ software. A minimum of 20 cell areas were measured for at least 3 independent experiments.

### D. Polyacrylamide Gel Preparation

Gels are prepared as described by Charrier et al. [16]. Fully polymerized chains of linear polyacrylamide (PAA) are enmeshed in crosslinked PAA networks to create viscoelastic gels containing separate elastic and viscous components. The linear PAA chains do not bond to the elastic network, and their effect is to increase the viscous contributions to the gel by allowing reorganization of the viscoelastic network via linear polymer relaxations. Chains of linear PAA were polymerized by making a solution 5.0% acrylamide in distilled water. This solution was degassed, then 0.024% ammonium per sulfate and 0.050% tetramethylethylenediamine (TEMED) were added to the solution. The linear PAA was allowed to polymerize for 2 hours at 37° Celsius. The elastic component of the polyacrylamide gel is synthesized by mixing acrylamide, bis-acrylamide, linear PAA, and distilled water. Polymerization was initiated by the addition of ammonium per-sulfate, and TEMED. Concentrations of elastic and viscoelastic gel components are listed in Table I.

**TABLE I.**
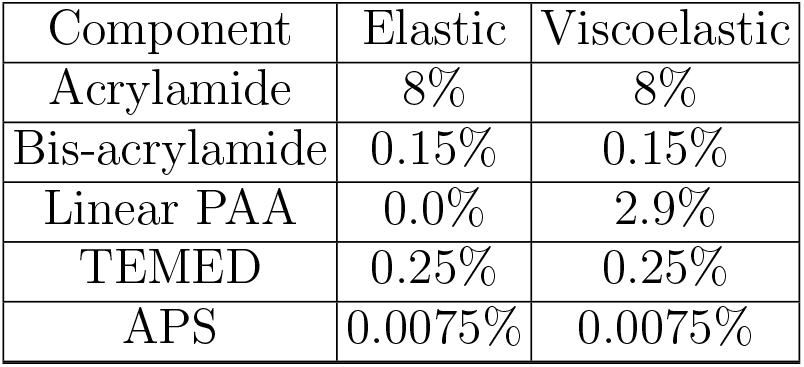
Recipes for elastic and viscoelastic gels, for gels treated with NHS 10% of distilled water is substituted for 0.4% NHS.

To facilitate cell adhesion with the substrate, we covalently link the PAA network with collagen I. The gels are designed so that collagen is bond to only the elastic component of the gel network by incorporating 0.04% NHS into its polymerizing solution. Gels were then incubated in a 50 *μ*g/mL collagen solution for 2 hr at room temperature. Then, gels were washed three times with PBS and sterilized by exposure to ultraviolet light.

### E. Rheological Measurement

Rheology measurements were performed on a Malvern Panalytical Kinexus Ultra+ rheometer (Malvern Panalytical, Westborough, Masachussets) using a 20 mm diameter parallel plate geometry. The elastic and viscoelastic gel solutions are polymerized at room temperatures between the rheometer plates at a gap height of 1 mm. The time evolution of polymerization is monitored by applying a small oscillatory shear strain of 2% at a frequency of 1 rad/sec for 30 minutes. The values for the elastic shear modulus and viscous shear modulus of each gel are determined by their plateau value once the gel has fully polymerized. Stress relaxation measurements were performed using a full polymerized sample by applying a constant shear strain of 5% and tracking the resulting stress relation with time.

### F. Statistical Analysis

Data presented as mean value ± standard errors (SE). Each experiment was performed at least twice. The unpaired Student’s t-test with two tails at the 95% confidence interval was used to determine statistical significance. Denotations: *, *p* <= 0.05; **, *p* <= 0.01; ***, *p* < 0.001; ns, *p* > 0.05.

## III. RESULTS

### A. Vimentin mediates cell spreading on viscoelastic substrates

To examine the effects of substrate viscous dissipation on cell spreading, we prepared polyacrylamide (PAA) hydrogels with elastic and viscoelastic material properties. PAA gels are a common model system for soft cell culture substrates. Once polymerized, acrylamide and bis-acrylamide form a linearly elastic network with time-independent responses to stress. To form a viscoelastic hydrogel, a dissipative element of linear PAA chains is incorporated into the network, as recently described in Charrier et al. [16]. A schematic of the viscoelastic gels is shown in Fig. 1a, where the linear PAA chains (pink) are integrated into the elastic crosslinked PAA network (cyan and green). The dissipative component of these gels relaxes in a time dependent response to an applied stress, imparting viscoelastic behavior to the gel.

**FIG. 1.**
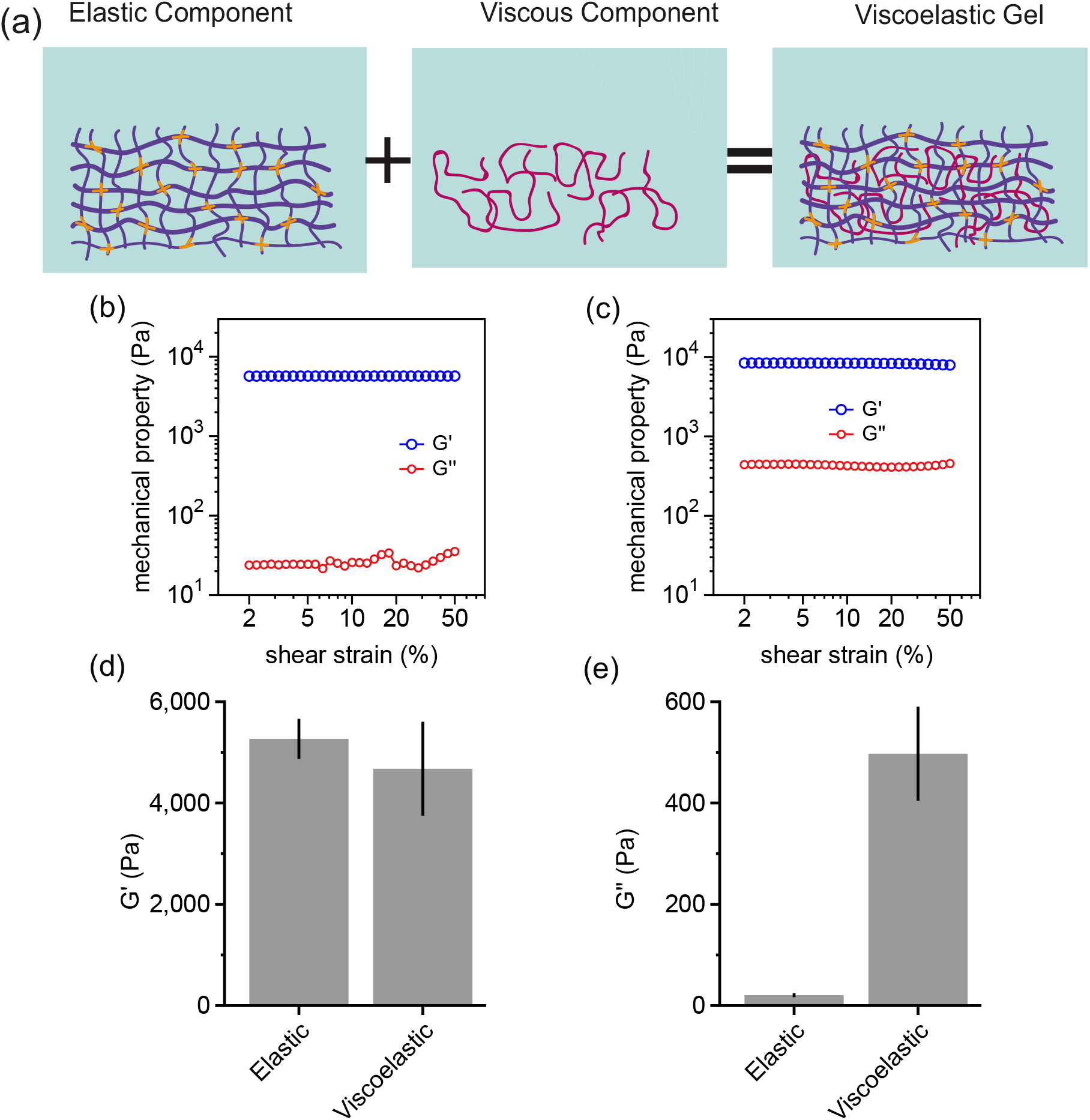
Characterization of viscoelastic polyacrylamide gels. (a) Schematic illustrating the elastic and viscous components of the viscoelastic gel. Acrylamide, cross-linker, and linear PAA chains are shown in blue, orange, and pink, respectively. (b) Shear strain sweep of elastic and (c) and viscoelastic hydrogels. The storage modulus G’ is approximately the same between the two gel types and is constant over a large of shear strain values. The loss modulus G” is nearly zero (within experimental noise) as expected in the elastic gel; while G” is finite and constant over varying shear strain in the viscoelastic gel. In (d) and (e) the average elastic shear modulus (G’) and viscous shear modulus (G”) at 2% strain are shown for both elastic and viscoelastic gels, respectively. N ≥ 3 gels per conditions, error bars denote standard error.

The mechanical properties of the gels are characterized by a shear storage elastic modulus *G*′ and a viscous loss modulus *G*″ via oscillatory rheology. To determine the effects of viscous dissipation independently of substrate stiffness on cell spreading, the gels were designed with a fixed storage modulus (*G*′ ≅ 5 kPa) but variable loss modulus *G*″ for the elastic (*G*″ ≅ 0 Pa) and viscoelastic gels (*G*″ ≅ 500 Pa), as measured at a frequency of 1 rad/s and 2% shear strain amplitude (Fig. 1). The loss modulus of the viscoelastic gel is thus 10% of its elastic moduli, which is in the 10-20% range of real tissue! [16].

If the time scale of substrate relaxation is similar to the time scale of cellular mechanosensing, the substrate relaxation may provide an important feedback for cells attempting to spread on viscoelastic substrates. To determine the time dependence of force dissipation on viscoelastic substrates, we performed stress relaxation measurements (Fig. 1e). A shear strain of 5% is continuously applied to the gel and the resulting stress relaxation is measured with time t. The stress relaxes to a finite value above zero, indicating the viscoelastic gels behave as a viscoelastic solid. The decrease in stress can be captured by an exponential decay function,

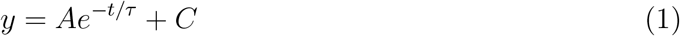

where *τ* is the characteristic timescales of force dissipation and A and C are fitting constants. By fitting this relationship to the data in Fig. 1e, we obtain a characteristic timescale *τ* of ≈ 14 seconds, which is expected to be relevant to cellular motion in the extracellular matrix environment [16].

**FIG. 2.**
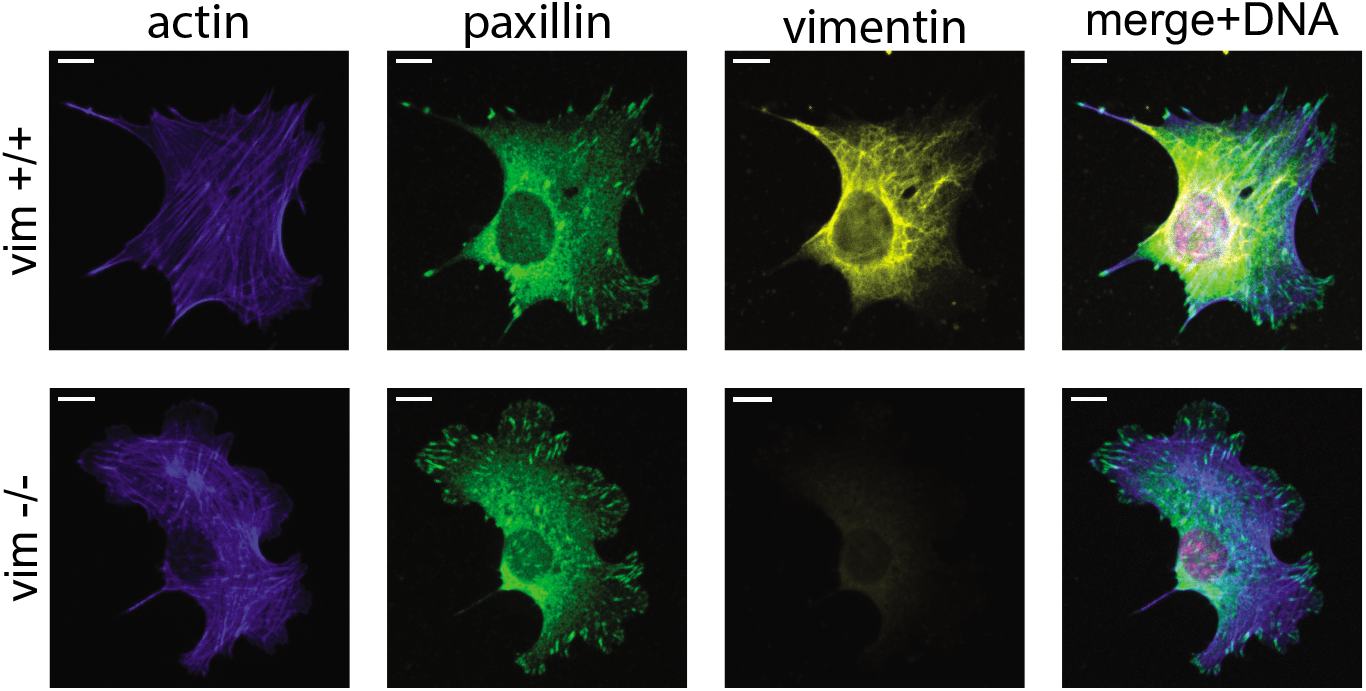
Representative images of wild-type (vim +/+) and vimentin-null (vim -/-) mouse embryonic fibroblasts (mEF) on glass slides. Wild-type and vimentin-null mEFs show similar distributions of F-actin stress fibers and paxillin focal adhesions. Confocal images show vimentin (yellow), actin (blue), paxillin (green) and the nucleus (pink). Scale bar, 10 *μ*m.

To determine the effects of vimentin to substrate viscous dissipation, we seeded wildtype and vimentin-null mouse embryonic fibroblasts (mEF) on the elastic and viscoelastic gels (Fig. 3a). To facilitate cells attachment to the substrates, the elastic component of the hydrogels was covalently linked with collagen I. Cells were imaged 24 hours after plating with bright field images used to quantify the cell spread area of single cells (Fig. 3b). We found that substrate viscoelasticity decreased cell spread area for both cell types, as seen on other viscoelastic hydrogels [16], but in a strongly vimentin-dependent manner. On purely elastic 5 kPa gels, the two cell types spread similarly (Fig. 3b) as on other soft elastic substrates [13]. However, on viscoelastic gels, their response was different (Fig. 3b): the vimentin-null mEF were significantly less spread compared to wild-type mEF. Taken together, these results indicate that vimentin contributes to how cells sense and respond to the physical properties of the extracellular matrix environment, particularly to substrate viscoelasticity.

**FIG. 3.**
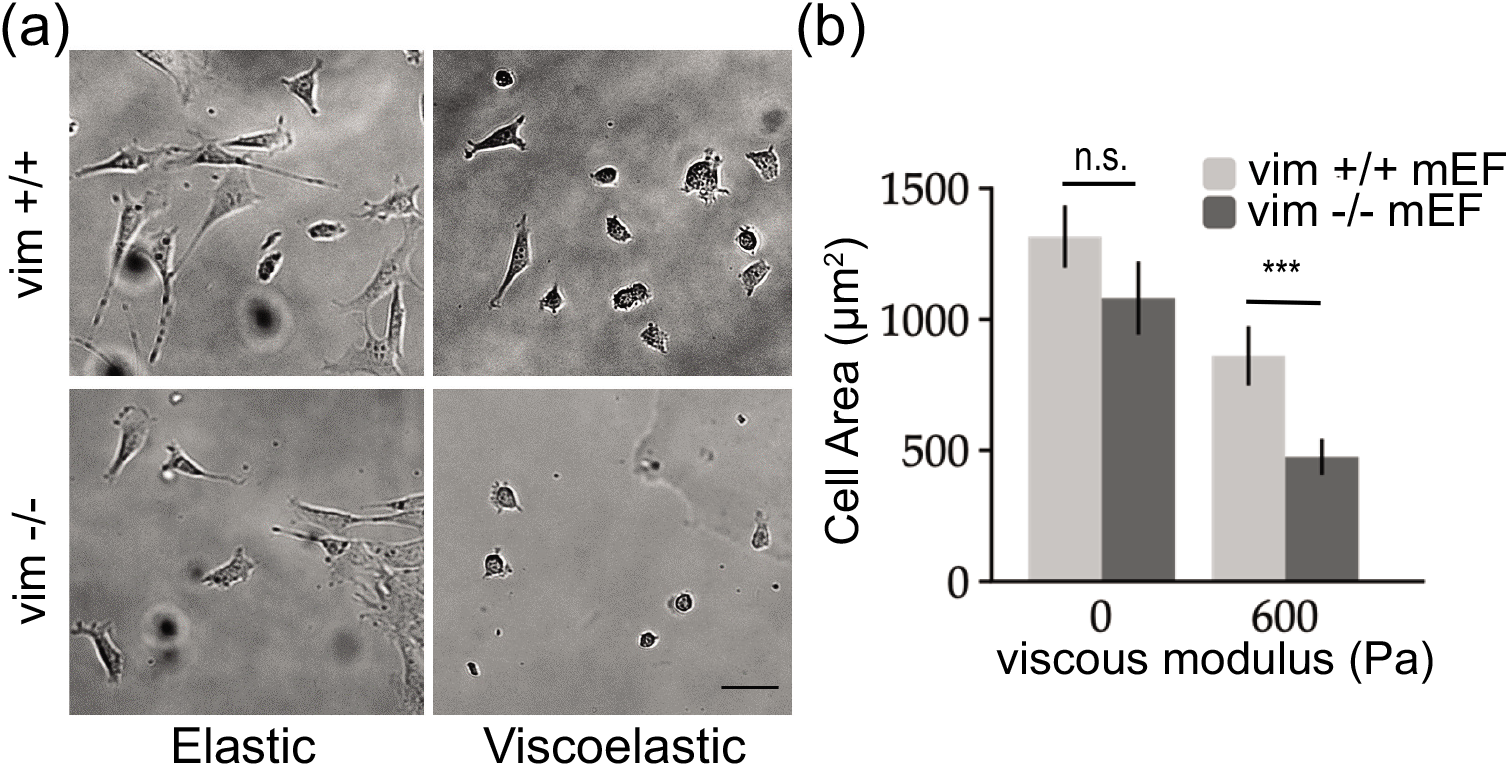
Vimentin enhances cell spreading on viscoelastic substrates. (a) Bright field images of mEFs after 24 hr of spreading on elastic and viscoelastic gels, coated with 50 *μ*g/mL collagen I. Scale bar = 30 *μ*m. (b) Average projected cell area of mEFs after 24 hr (n ≥ 20 cells over N ≥ 3 experiments per conditions. Error bars denote standard error).

### B. Effects of substrate viscoelasticity on cell-substrate and cell-cell interactions

We next examined how substrate viscous dissipation affects cell-substrate and cell-cell interactions (Fig. 4). To quantify cell-substrate adhesions, we counted the total number of adherent cells after 24 hr. On both elastic and viscoelastic substrates, wild-type cells were more successful in remaining adhered to the substrate when compared to vimentin-null cells. This observation was more striking on viscoelastic substrates, where it was seen that on average wild type cells were 3-fold more likely to remain adhered (Fig. 4a). These results suggest that cell adhesion to viscoelastic substrates correlates with vimentin expression.

**FIG. 4.**
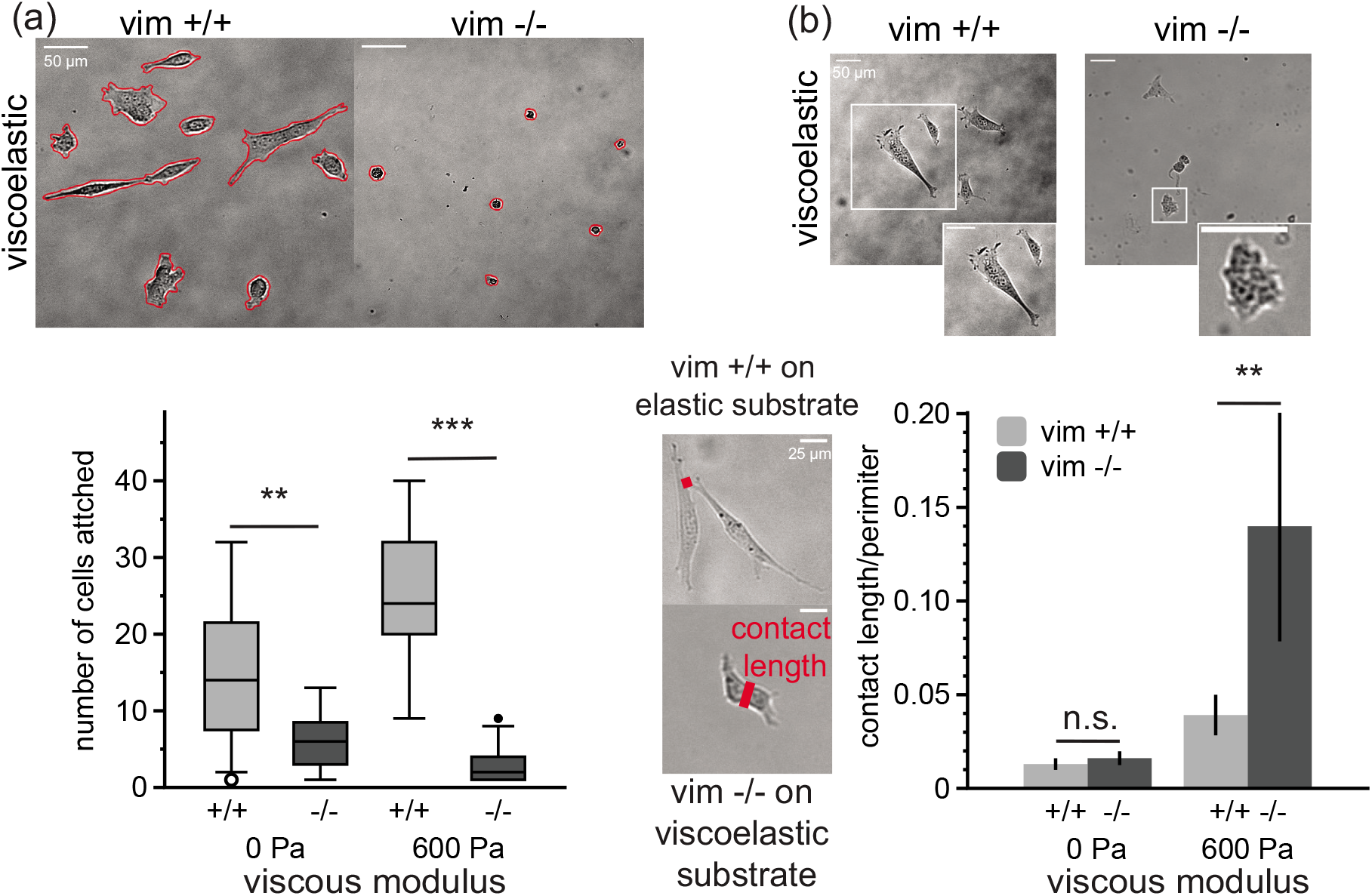
Cell-cell contacts facilitate cell attachment to substrates with viscous dissipation. Panel (a) shows the average number of cells attached on gel after 24 hours. Cells are counted inside of a 0.5 mm^2^ imaging window. (b) Quantification of cell-cell contacts. In vimentin-null cells, cells form greater contact with neighboring cells, as measured by the ratio of the cell contact length to the perimeter of neighboring cells. (Cells attached: n ≥ 10 images per condition, N ≥ 2 experiments. Contact length: n ≥ 15 cells per condition, N ≥ 2 experiment. Error bars denote standard error).

Cell spreading depends on the mechanical properties of the substrate matrix but also direct contact-mediated cell-cell interactions. We observed different rates of cell clustering behavior between our wild-type and vimentin-null mEF on viscoelastic substrates. Vimentin-null mEF adhered to viscoelastic substrates formed noticeable more clusters after 24 hours; whereas cells with vimentin or on elastic substrates did not cluster tightly. To assess cell-cell interactions, we quantified cell-cell interactions by the length of direct contact between two touching cells, which is independent of the adherent cell density. Fig. 4b shows two examples of these cell-cell interactions. For wild-type cells on elastic substrates, cell-cell contacts are small compared to the cell’s contact with the substrate, whereas the vimentin-null cells on viscoelastic substrates are strongly coupled. To normalize the cell-cell contact with cell spread area, we define the cell-cell contact by dividing the contact length by the total perimeter of the pair of cells. As shown in Fig. 4, we find that substrate viscoelasticity increases cell-cell contact for both cell types, indicating that cell-cell interactions promote adhesion on viscoelastic substrates. While cell-cell contact is similar between wild-type and vimentin-null meF on elastic substrates, cell-cell contact increases more than 200% for vimentin-null cells compared to wild-type cells on the viscoelastic substrates. Taken together, these results indicate that vimentin may facilitate adhesion to viscoelastic substrates by promoting cell-matrix adhesions over cell-cell interactions.

### C. Substrate viscoelasticity mediates vimentin organization

Next, we analyzed the organization of vimentin networks on elastic and viscoelastic substrates. Vimentin networks are apparent in wild-type mEF on all substrates but the organization of the vimentin networks varied, as shown in immunofluorescence images in Fig. 5. On the elastic substrates, vimentin was spread throughout the cytoplasm of the cells, with filamentous bundles extending toward the periphery of the cells. Filamentous vimentin bundles such as these have been implicated in load-bearing units in the cell, contributing to proper alignment of actin-based cell traction stress [22]. However, on viscoelastic substrates, the vimentin network was more condensed in a mesh-like cage around the cell nucleus and vimentin filamentous strands were less evident.

**FIG. 5.**
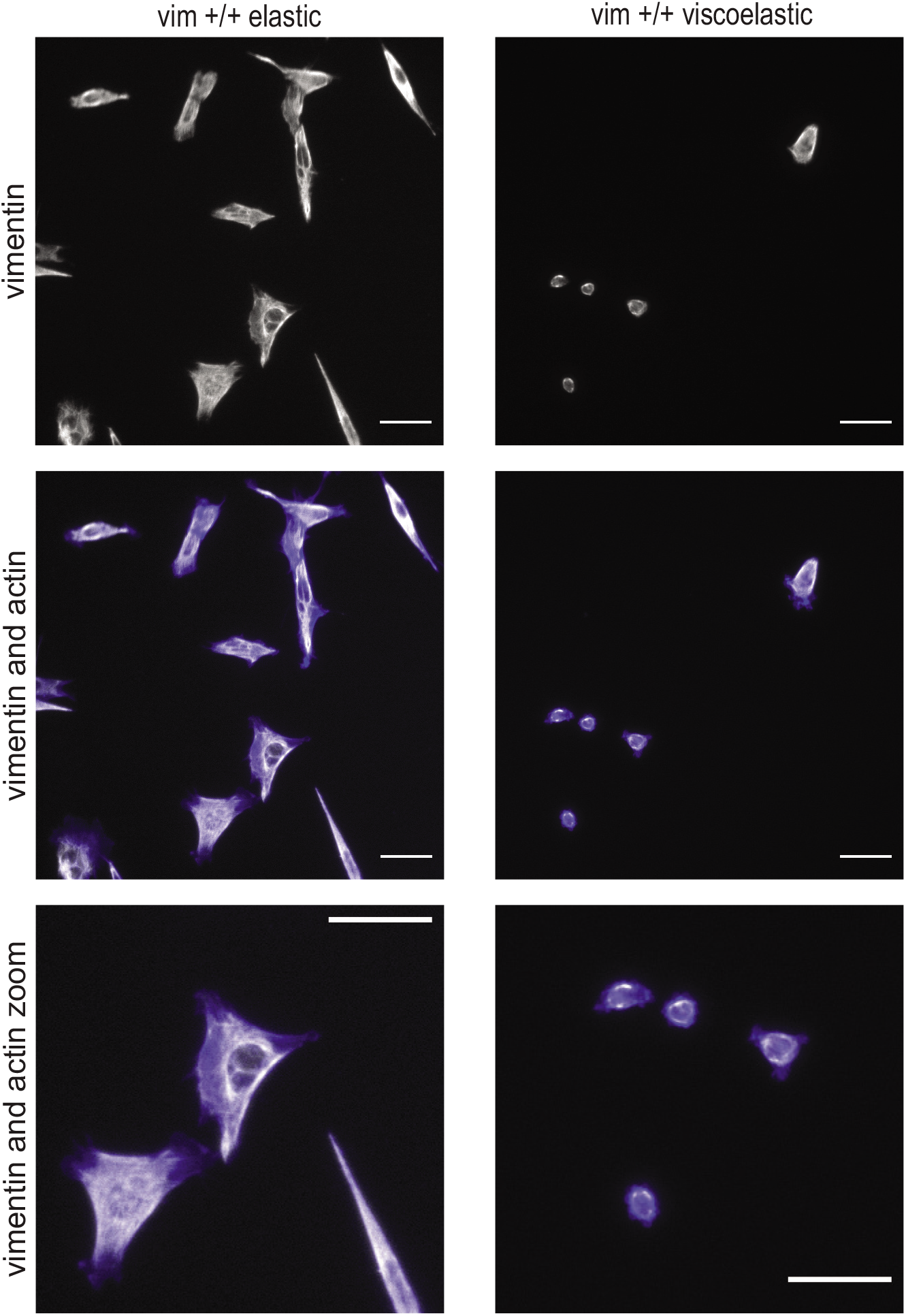
Viscoelastic substrates decrease cell spreading and alter vimentin organization. Representative immunofluorescence images of wild-type cells on elastic and viscoelastic gels. On elastic gels, the vimentin cytoskeletal network extends out towards the periphery of the cell; while on viscoelastic gels, the vimentin network is more collapsed around the nucleus. Vimentin is shown in grey and actin is shown in blue. Scale bar, 50 *μ*m.

When cultured on viscoelastic gels, the fibroblasts were less spread, suggesting a reduced ability to form stable attachments to substrates. Cells form adhesion sites with the substrate though integrins and the extracellular matrix ligands to which they bind. To visualize cell-matrix adhesions, we fixed cells and stained for antibodies against paxillin, a cellular protein found in cell focal adhesions sites (Fig. 6). Most of the cells (>70%) displayed paxillin patches and actin stress fibers on elastic substrates. This was reduced by nearly 50% on viscoelastic gels in both wild-type and vimentin-null mEF. These results suggest that substrate viscoelasticity alters focal adhesion formation and actin architecture. The number of cells with focal adhesions and stress fibers were similar between wild-type and vimentin-null cells, suggesting an independent effect of vimentin in enhancing cell spreading on viscoelastic substrates.

**FIG. 6.**
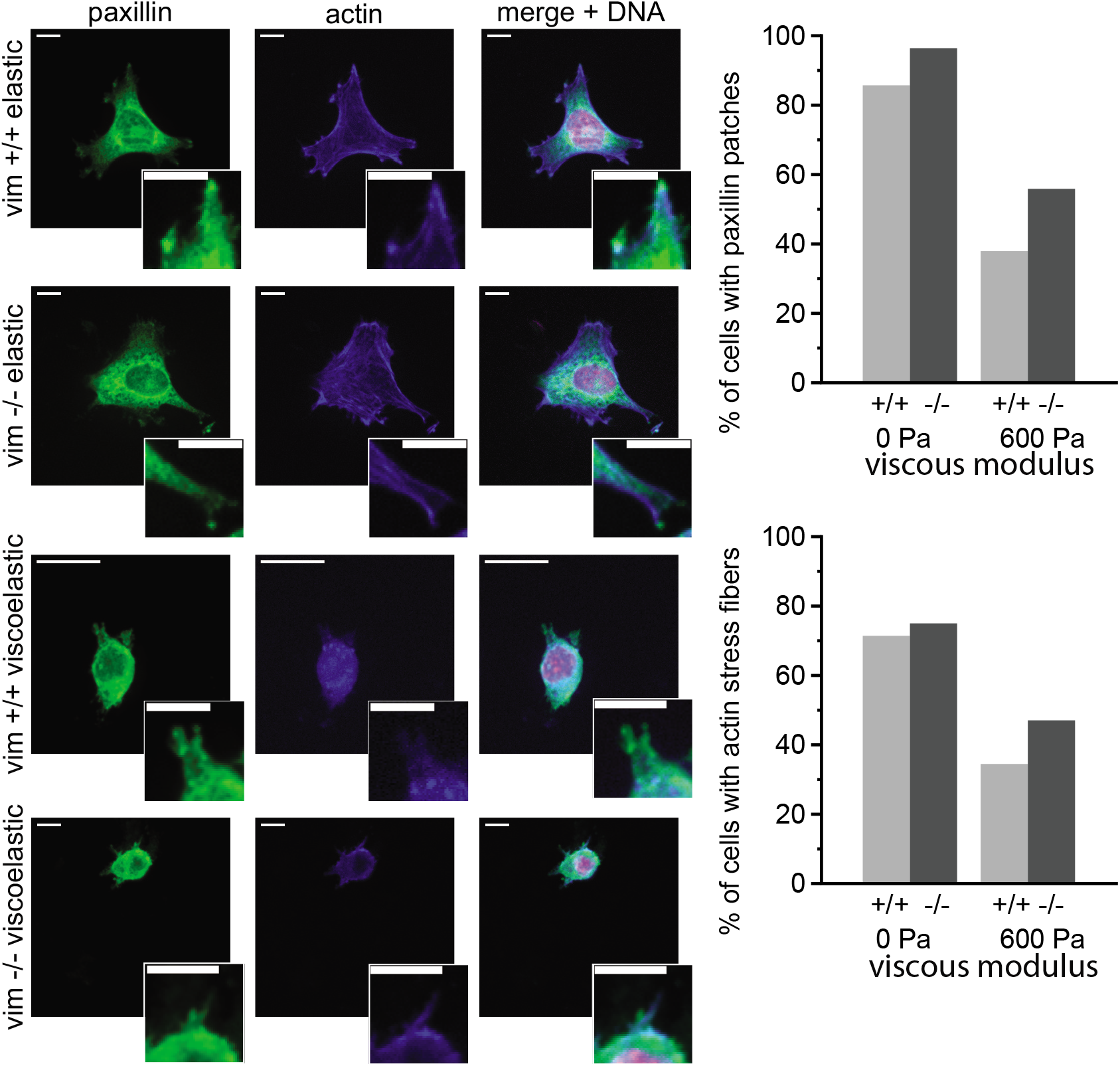
Distribution of actin stress fibers and paxillin focal adhesions on elastic and viscoelastic substrates. Immunofluoresence images of actin (blue), paxillin (green), and DNA (pink) on elastic and viscoelastic substrates. Panel (a) shows spinning disk confocal images of both wild type (vim +/+) and vimentin-null (vim -/-) mEF that have been incubated for 24 hours on elastic and viscoelastic substrates coated with collagen I. Scale bar, 10 *μ*m. Panels (b) and (c) present the percentage of cells that exhibit actin stress fibers and paxillin patches, respectively. (n ≥ 20 cells per condition, N = 2 experiments, error bars denote standard error).

## IV. DISCUSSION

Cell spreading is a complex process that probes fundamental interactions between the behavior of the cell and its coupling to the underlying substrate matrix. This process depends on the cellular cytoskeleton and focal adhesion complexes that attach the cell to the substrate. The intermediate filament protein vimentin contributes to cell morphology and the mechanical resilience of cells, but its role in how cells sense and respond to physical properties of the environment is largely unclear. Here, we provide evidence that vimentin is an integral element of cellular mechanical sensing, particularly on substrates with viscous dissipation.

We found that wild-type (vim +/+) and vimentin-null (vim -/-) mEFs spread similarly on soft purely elastic substrates, in agreement with prior experiments [9, 20]; however, cells lacking vimentin have decreased spreading on viscoelastic substrates. Substrate viscoelasticity altered the vimentin network structure, collapsing filamentous vimentin bundles into a mesh-like cage around the nucleus. Based on models of vimentin as a load-bearing structures that helps align actin-based stresses [22], this observed change in vimentin organization suggests that vimentin is under less mechanical tension on viscoelastic substrates compared to elastic substrates. On viscoelastic gels, cells are less spread and have fewer paxillin patches, indicating a weaker cell-substrate interaction. Loss of vimentin correlated with a fewer number of substrate adherent cells and a preference for cell-cell interactions (Fig. 4).

Cells are thought to sense their substrate through a “motor-clutch” mechanism [23]. In this context, focal adhesions act as molecular clutches, physically linking the cellular cytoskeleton to the extracellular matrix substrate. These physical linkages transmit traction forces to the underlying substrates. Motor clutches are generally assumed to act as slip bonds, with a dissociation rate that increases with the amount of force it bears by coupling the cell with the substrate. As motors bind with the substrate, they generate friction and resist the retrograde flow of F-actin filaments. This allows actin polymerization to push the leading edge of the cell forward, which ultimately results in cell spreading.

The strong effect of substrate viscoelasticity on vimentin-dependent cell spreading might be surprising given that vimentin is dispensable for cell spreading on soft elastic substrates [13]. On the other hand, filamentous vimentin networks are less dynamic [24] and more viscoelastic [25] compared to the actin and microtubule networks in the cytoskeleton. Thus, the stabilizing effect of vimentin on microtubule orientation [26] and actin-based stresses [22, 27] may be particularly evident for cell spreading on viscoelastic substrates, which act to dissipate and effectively lessen cell-generated traction stresses.

On viscoelastic substrates, the dynamics of focal adhesion assembly and disassembly compete with substrate viscous dissipation to facilitate cell spreading. Recent motor-clutch models [17] have been developed to capture the effects of adhesion dynamics and substrate viscoelasticity on cell spreading. On viscoelastic substrates, cells feel a time-dependent effective stiffness that decreases over a characteristic substrate relaxation time scale. This results in weaker adhesions and faster retrograde flows, which decrease cell spreading. Prior experiments using Fluorescence Recovery After Photobleaching (FRAP) analysis of GFP-paxillin showed that the rate of paxillin turnover in MCF-7 cells is significantly increased in cell with high levels of vimentin expression [20]. If vimentin increases the binding time of the clutches, then clutches can quickly bind and break without forming large stable focal adhesions. The time scale of binding is larger than the lifetime of a focal adhesion. This results in an effective “frictional slippage” regime, which stalls retrograde flow and promotes cell spreading. Since focal adhesion turnover is slower in cells lacking vimentin, they are predicted to sense a lower effective substrate stiffness. In this case, the clutch binding time is less than the lifetime of the motor clutches, resulting in a “load and fail” regime. The adhesive forces are expected to be smaller than the frictional slippage regime in vimentin expressing cells, resulting in increased retrograde flow and reduced cell spreading. Overall, our results suggest that vimentin could promote cell spreading on viscoelastic substrates by mediating emergent interactions between focal adhesion dynamics and substrate relaxation time scales.

Taken together, our results indicate a new function of vimentin intermediate filaments in modulating cellular response to viscoelastic environments that has important implications for understanding cell and tissue functions. Intermediate filaments play diverse roles in a range of cell and tissue functions [28, 29] and are important to maintaining cell morphology and adhesion [18]. Vimentin IF in particular has been implicated in cataracts [30], coronaviruses [31], wound healing [25], and many forms of metastatic cancer [32–34]. Our results indicate that vimentin promotes cellular adhesion and motility through viscoelastic environments, such as extracellular matrices and in vivo tissue. Impaired wound healing and cell migration in cells lacking vimentin could result from a reduced ability for cells to interact strongly with viscoelastic tissue. More broadly, our results show that vimentin contributes to how cells respond to viscoelastic properties of the extracellular matrix where applied stresses dissipate on timescales relevant to cellular mechanical sensing

## V. ACKNOWLEDGEMENTS

We acknowledge useful discussions with Jennifer Schwarz, Vivek Shenoy, Farid Alisafaei, Bobby Carroll, and Paul Janmey. This work was supported by the National Science Foundation MCB 2032861 awarded to A.E. Patteson.

